# The functional brain organization of an individual predicts measures of social abilities in autism spectrum disorder:Predicting symptoms in autism with brain imaging

**DOI:** 10.1101/290320

**Authors:** Evelyn MR Lake, Emily S Finn, Stephanie M Noble, Tamara Vanderwal, Xilin Shen, Monica D Rosenberg, Marisa N Spann, Marvin M Chun, Dustin Scheinost, R Todd Constable

## Abstract

Autism Spectrum Disorder (ASD) is associated with multiple complex abnormalities in functional brain connectivity measured with functional magnetic resonance imaging (fMRI). Despite much research in this area, to date, neuroimaging-based models are not able to characterize individuals with ASD with sufficient sensitivity and specificity; this is likely due to the heterogeneity and complexity of this disorder. Here we apply a data-driven subject-level approach, connectome-based predictive modeling, to resting-state fMRI data from a set of individuals from the Autism Brain Imaging Data Exchange. Using leave-one-subject-out and split-half analyses, we define two functional connectivity networks that predict continuous scores on the Social Responsiveness Scale (SRS) and Autism Diagnostic Observation Schedule (ADOS) and confirm that these networks generalize to novel subjects. Notably, these networks were found to share minimal anatomical overlap. Further, our results generalize to individuals for whom SRS/ADOS scores are unavailable, predicting worse scores for ASD than typically developing individuals. In addition, predicted SRS scores for individuals with attention-deficit/hyperactivity disorder (ADHD) from the ADHD-200 Consortium are linked to ADHD symptoms, supporting the hypothesis that the functional brain organization changes relevant to ASD severity share a component associated with attention. Finally, we explore the membership of predictive connections within conventional (atlas-based) functional networks. In summary, our results suggest that an individual’s functional connectivity profile contains information that supports dimensional, non-binary classification in ASD, aligning with the goals of precision medicine and individual-level diagnosis.

## INTRODUCTION

The global prevalence of autism spectrum disorder (ASD) in 2010 was estimated to be 7.6 per 1000, or approximately 52 million cases.^1^ The lifetime burden of ASD is greater than that of both attention deficit hyperactivity disorder (ADHD) and conduct disorders combined, totalling 6.2 million disability-adjusted life-years globally.^1, 2^ Diagnosis of ASD continues to be challenging, particularly in young children, in part because ASD includes a wide range (or spectrum) of symptoms, skills, and levels of impairment.^3^ As such, specific diagnoses, assessments of symptom severity, and choice of treatment for each symptom domain often varies widely across individuals. Reflecting this clinical complexity, the associated neural correlates of ASD are also complex, have been difficult to characterize, and are not well understood.^4^

Magnetic resonance imaging (MRI) has been used to discover structural and functional differences between ASD and typically developing (TD) individuals.^4–7^ Functional MRI (fMRI) is a non-invasive imaging methodology that can measure a correlate of brain activity reflected by changes in local tissue oxygenation.^8^ The dynamic time-series fMRI data can be analyzed to identify patterns of coupling between distinct anatomical regions [^9^]: a measure referred to as functional connectivity. A map of all the connections in the brain is referred to as the functional connectome.^10^ Recent work has demonstrated that individual’s have unique functional connectivity patterns, that contain information about behavioral traits and/or clinical symptoms.^11–13^ Such connectome-based assessment of an individual’s brain organization may prove useful in guiding the clinical management of patients.^11, 14–16^

Functional connectivity studies of ASD have shown alterations in multiple functional networks compared to typically developing (TD) individuals.^4, 5, 7, 17,-30^ However, studies demonstrating a continuous relationship between behavioral measures (gold-standard clinical evaluation) and connectivity are limited, many studies are under-powered, and the results are rarely replicated.^17, 31^ Furthermore, very few studies predict out-of-sample — rather than explain within-sample — clinical scores.^32, 33^ As recent reviews have summarized, findings in this area are generally complex and non-converging, likely reflecting both the daunting heterogeneity of ASD and the disparate but relevant brain circuits investigated.^34–36^

Given the substantial individual differences in ASD symptomatology and the complex imaging correlates, a whole-brain data-driven dimensional approach focused on individual differences rather than categorical/binary grouping may be more useful to capture the subtle features that involve the interplay of multiple brain regions. In this work we test the hypothesis that connectome-based predictive modeling (CPM) can be used to identify complex whole-brain networks that predict symptom severity based only on an individuals’ functional connectome.^11, 16, 37^ We focus on two clinical scores relevant to autism, the Social Responsiveness Scale (SRS) and the Autism Diagnostic Observation Schedule (ADOS), available from the Autism Brain Imaging Data Exchange (ABIDE) consortium.^38, 39^ Using both leave-one-subject-out (LOO) and split-half cross-validation (CV), we validate these models and identify two anatomically distinct functional networks related to SRS and ADOS scores. Finally, motivated by the overlap in symptomatology and genealogy between ASD and attention-deficit/hyperactivity disorder (ADHD), as well as the high co-occurrence of these disorders within individuals, we explore the generalizability of our SRS and ADOS models in an independent data set derived from the ADHD-200 Consortium.^40^

This dimensional rather than binary categorical approach captures degree of severity, which is of particular importance in ASD where a broad range of phenotypes are a salient feature of the disorder. Furthermore, CPM preserves the ability to track response to therapies and captures a range of clinical presentations, including (a)symptomatic siblings of ASD individuals. It is also in line with the National Institute of Mental Health’s conceptualization of mental health disorders.^41^ In summary, we demonstrate that behavioral measures and imaging data can be used to develop models relating connectivity to symptom severity in a dimensional approach at the individual subject level.

## METHODS

### 2.0 Data sets

We analyzed data from ABIDE-I/II and the ADHD-200 consortium, two publicly available multi-site data sets of resting state fMRI (rs-fMRI), demographic, and clinical assessment data.^38–40^ Detailed information is available for ABIDE-I/II at fcon_1000.projects.nitrc.org/indi/abide/ and ADHD-200 at fcon_1000.projects.nitrc.org/indi/adhd200/. Refer to supplementary material for an imaging parameter summary.

### 2.1 Rs-fMRI data processing

Standard pre-processing procedures were used as previously described.^12^ Motion correction was performed using SPM8 (fil.ion.ucl.ac.uk/spm/). Images were iteratively smoothed to a full-width half maximum of 6mm to reduce motion related confounds.^42^ All further analyses were performed using BioImage Suite.^43^ Covariates of no interest were regressed from the data including: linear and quadratic drifts, and mean cerebral-spinal-fluid, white and gray matter signals. For additional control of motion related confounds, a 24-parameter motion model (including six rigid-body motion parameters, six temporal derivatives, and these terms squared) were also regressed from the data. Frame-to-frame motion was estimated as the Euclidean distance between the center of gravity of neighboring frames from the transformation matrix, which incorporated three translation and three rotation estimates. We applied temporal smoothing with a Gaussian filter (cutoff frequency=0.12Hz).

Each individual’s functional connectome was calculated using a functionally defined atlas of 268 cortical and subcortical nodes defined in a separate population.^11, 44^ For each subject, the atlas was warped from MNI space into single-subject space via concatenation of a series of linear and nonlinear registrations as previously described.[^12^] All transformation pairs were calculated independently, combined into a single transform, and inverted, warping the functional atlas into single participant space. For each individual, a 268x268 connectivity matrix was calculated using Pearson

### 2.2 Behavior metrics

**ABIDE-I/II.** SRS serves as a broad-spectrum estimate of autistic traits. It is commonly applied not only in individuals with ASD, but also in family members of ASD individuals and trans-diagnostically. ADOS is more strictly applied to assess and diagnose individuals with ASD. Briefly, SRS is a 65 item questionnaire. Individuals are scored out of four: (1) ‘not true’, to (4) ‘almost always true’, by either themselves, or a parent/guardian.^45–47^ The following is an example question from SRS: “avoids starting social interactions with others”. ADOS is more intensive, and based on the observation/evaluation of elicited imaginative activities involving social role-playing and communication within a standardized context (e.g. telling a story) by a trained observer.^48^ For both scales, a greater score indicates a deficiency in reciprocal social behaviors and a likelihood that the individual will find everyday social interactions challenging. Six SRS and eight ADOS sub-scale scores (from two modules) were available. SRS and ADOS sub-scale scores are highly correlated within scales (i.e., SRS sub-scale scores with other SRS sub-scale scores, and ADOS sub-scale scores with other ADOS sub-scale scores) but generally less correlated between scales (i.e., SRS sub-scale scores with ADOS sub-scale scores) [Supplementary Figure 1.A.].

**ADHD-200.** We include ADHD symptom measures from the ADHD Rating Scale-IV.^49^ These scores are calculated by summing responses to 18 questions on a 4-point scale: (0) ’rarely or never’, to (3) ’always or very often’. Questions assess attention (e.g. “is easily distracted by extraneous stimuli”) or hyperactivity (e.g. “interrupts or intrudes on others”).

### 2.3 Model building: behavior prediction

Models were built using LOO-CV CPM analysis as described previously.^11, 12, 37^ Briefly, an iterative three-step analysis was performed: (1) feature selection (N-1 training set), (2) building of a predictive model (N-1 training set), and (3) testing on the left-out subject. Each individual was left-out of the training set once in this iterative framework. In the first step, across individuals in the training set, a Pearson correlation is calculated between each functional connection, or edge, of the 268x268 connectivity matrix and clinical score. The resulting set of correlations is thresholded (P<0.01) to create feature sets that correlate either positively (+ve) or negatively (-ve) with the clinical measure. In the second step, ‘network strength’ (a single number reflecting the sum of all edges in the feature set) is computed for each individual in the training set. Network strength is a subject-specific summary statistic akin to a weighted degree.^50^ Next, linear regression is used to build a model of the relationship between the clinical score and network strength across individuals. Finally, this linear model along with the network strength from the left-out subject is used to predict the clinical score for the left-out individual. The predictive power of the model is assessed by the Pearson correlation of predicted versus measured behavioral score across all individuals. We apply Bonferroni correction for multiple comparisons (6 SRS and 4 ADOS scores).

### 2.4 Internal validation: (1) split-half CV, (2) permutation testing, and (3) extrapolation

To test model robustness, we use (1) split-half validation (n=200 iterations) and (2) permutation testing (n=1,000 iterations). For split-half validation, individuals are divided equally between train and test groups by random selection. Network/model building is conducted within the training group and the model applied to the test group. Permutation testing was conducted as described previously.[^37^] Briefly, subject labels and clinical scores were randomly shuffled to break the true brain-behavior relationship, then prediction (LOO-CV) performed on the shuffled data to generate a null result. We test if correlations from train/test and shuffled data come from different distributions (*kruskalwallis,* MATLAB). As a this validation step (3), for each split-half iteration, networks/models were applied to all individuals (less those used to generate the model) whether or not clinical scores were available from these individuals (N=ABIDE-I/II-training). Thus, for each individual in ABIDE-I/II, we generate clinical score predictions which we compare between ASD and TD groups and demonstrate across this larger sample that the scores differ significantly (*kruskalwallis*, MATLAB). We apply Bonferroni correction for multiple comparisons (6 SRS and 4 ADOS scores).

### 2.5 Network anatomy

The brain is complex and the networks identified by CPM reflect this complexity. In order to assess the extent to which the SRS and ADOS models share common features, we compute the probability that *n* shared edges exist between SRS/ADOS networks and edges within or between 10 *a priori* defined atlas networks.[^44, 51^] Significance was determined using the hypergeometric cumulative distribution function (*hygecdf*, MATLAB, Bonferroni correction for 55 comparisons). We report the likelihood (1.0-Pvalue) that each atlas network (and inter-network pair) contributes to SRS and ADOS networks. Furthermore, we analyze the distribution of edge lengths (defined as the Euclidean distance between the center of mass between each node) within networks. Note that this estimate of geometric distance is a rough proxy for synaptic distance. Using MATLAB, we test for outliers (*kurtosis*), normalcy (*lillietest*), a tendency towards long/short connections (*skewness*) and differences between +ve/-ve network distributions (*ranksum*). In addition, we evaluate the anatomy of shared features between networks by taking the products of +ve/+ve, +ve/-ve, -ve/+ve and -ve/-ve network pairs across scales and compute the likelihood that each of the resulting sets of shared features contain *n* edges from atlas-networks.

## RESULTS

### 3.0 Participants

Due to the sensitivity of functional connectivity measures to motion, we select subjects with frame-to-frame motion <0.08mm.^52^ Furthermore, we exclude individuals without sex or age information, thereby reducing ABIDE-I/II to 632 individuals (N=290/342, ASD/TD). Individuals included in each of our analysis steps are illustrated in Supplementary Figure 1.B. Clinical scores were independent of sex and medication status [Supplementary Figure 1.C./D.]; therefore, individuals were not excluded according to either category. Where reported (ABIDE-II), individuals with eyes closed were excluded. Full intelligence quotient (FIQ), age, and motion were limited such that known and predicted clinical scores were independent of these nuisance variables. Thus, SRS and ADOS groups were reduced to N=260/352 and 58/79. The male/female, and TD/ASD ratios as well as mean ± SD and range (min.-max.) of motion (mm), FIQ and age (yrs.) are summarized in Supplementary Figure 2. Data from the ADHD-200 consortium, collected at Peking University were thresholded according to the same methodology (N=77/35, ADHD/TD).

### 3.1 ASD behavior prediction (ABIDE-I/II)

For all SRS sub-scales, predicted behavior from LOO-CV analyses correlated with known scores (R=0.23-37, P<0.00002) [Figure 1.A.]. Similarly, for the majority of ADOS sub-scales, predicted clinical score correlated with known score (R=0.43-0.60, P<0.0002) [Figure 1.B.]. In all cases, age, FIQ and motion were included along with our model as covariates [Supplementary Table 1]. As a secondary analysis, we considered male individuals and obtained comparable results [Supplementary Table 1.C./D.]. Insufficient data was available for a female group. These results affirm our hypothesis that connectome based predictive modeling can be used to predict severity of ASD clinical symptoms. That is, the individual’s functional connectome contains information reflecting social behavioral scores as measured by SRS and ADOS.

**Figure 1.**
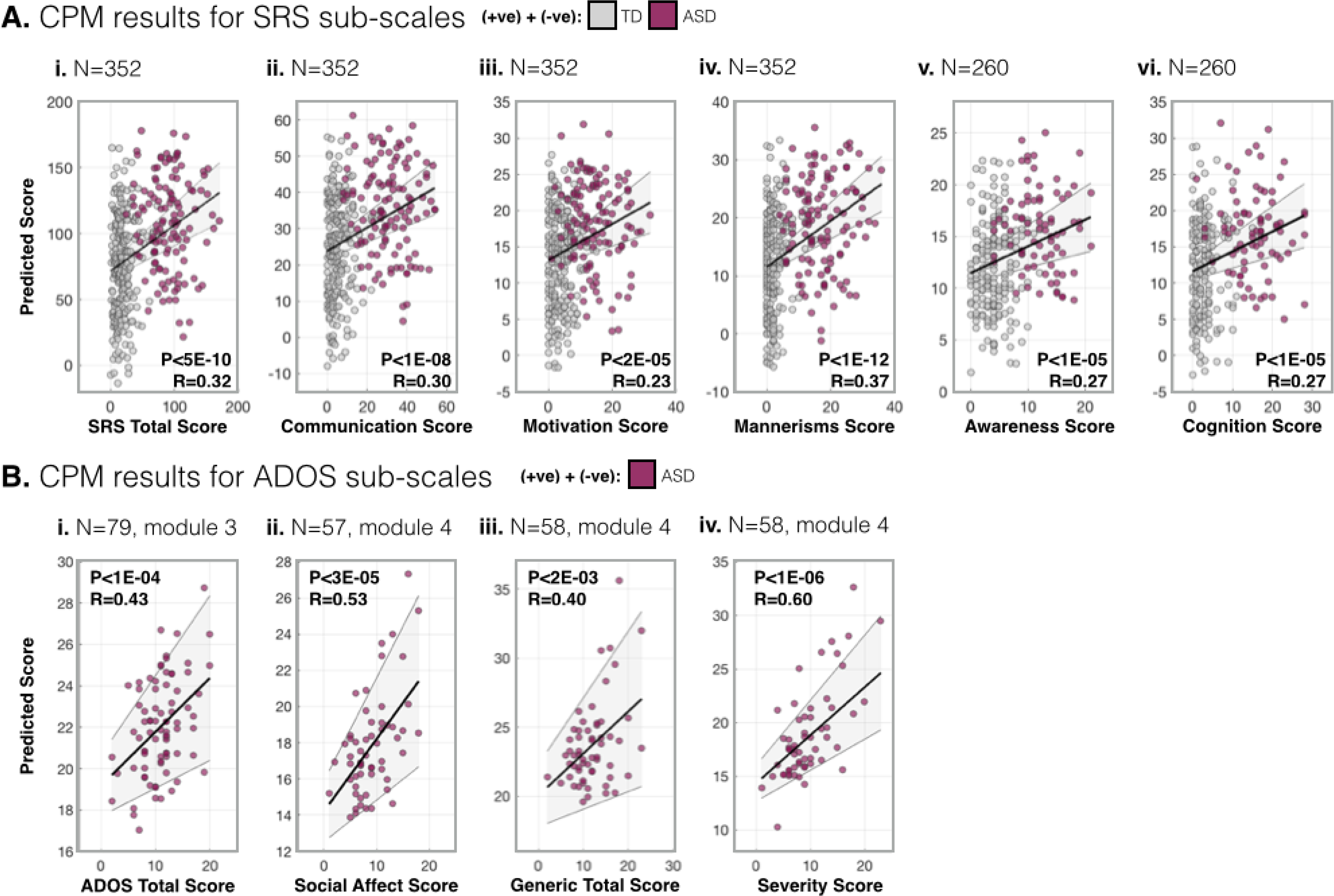
LOO-CV CPM results for SRS and ADOS sub-scale scores. (**A.**) LOO-CV CPM results for SRS sub-scale scores. For each SRS sub-scale **(i.-vi.)**, the sum of the predicted SRS score from +ve and -ve models are plotted against known scores. (**B.**) As in (A.) for ADOS sub-scale scores. The linear regression and 95% confidence interval are shown in black/grey.

### 3.2 Internal validation of SRS/ADOS models

To test the robustness of LOO-CV SRS/ADOS models and generalizability within the ABIDE-I/II, we used split-half CV and permutation testing. Correlations between known and predicted behavior for split-half train/test groups are plotted alongside correlations obtained from shuffled data (null results) [Figure 2.A.]. For all SRS and ADOS sub-scales, correlations from train/test data were greater than shuffled data. When SRS/ADOS models were applied to all individuals (less those used to generate the model), predicted scores were greater for ASD relative to TD individuals (N=632) [Figure 2.B] with the exception of the ADOS severity sub-scale. As a control, whole-brain connectivity in place of SRS/ADOS networks showed no difference between diagnostic groups [Supplementary Figures 3&4.C.]. Note, networks/models generated within this section are not used in future sections. Networks from Section 3.1 (generated from all individuals) are applied in all following analyses.

**Figure 2.**
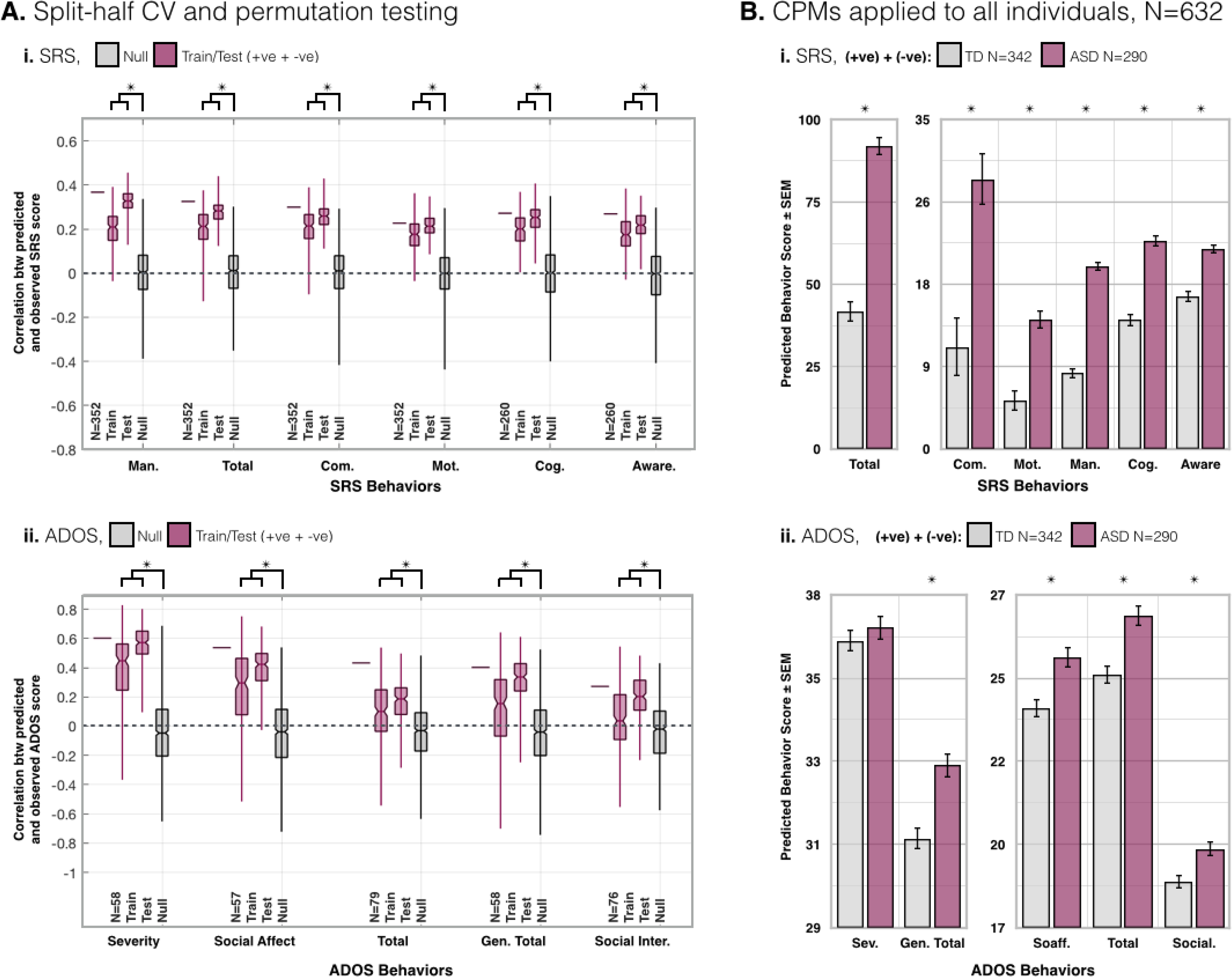
SRS and ADOS split-half CV and permutation testing results, and application to all individuals (ABIDE-I/I I) (**A**.) Correlation (R-value) for split-half CV. (**i.**) For each SRS sub-scale, the bar in the left-most column is reproduced for reference from the LOO-CV CPM results reported in Figure 1. N=352/260. The middle two columns are from split-half train/test CV CPMs (n=200 iterations). The final column shows the null results form permutation testing where subjects and scores are scrambled prior to LOO-CV (n=1,000). For all SRS sub-scales, train/test results are greater than null results (P<2E-144). (**ii.**) Same as (**i.**) for ADOS sub-scales (P<0.03). See Supplementary Figure 2.A.&3.A. for results from SRS&ADOS +ve/-ve feature sets. (**B.**) From each iteration of the split-half CV, the model was applied to all individuals from ABIDE-I/II less those in the training group (N=632-training) to predict clinical scores. Across iterations (n=200) mean predicted scores are compared between TD and ASD individuals. For all sub-scales, predicted SRS (P<1E-07) scores are greater for ASD than TD individuals (**i.**). Likewise, all but the severity ADOS sub-scale score was greater for ASD than TD individuals (P<0.02) (**ii.**). Between ASD and TD groups, motion (P>0.14), and age (P>0.96) were not different.

**Figure 3.**
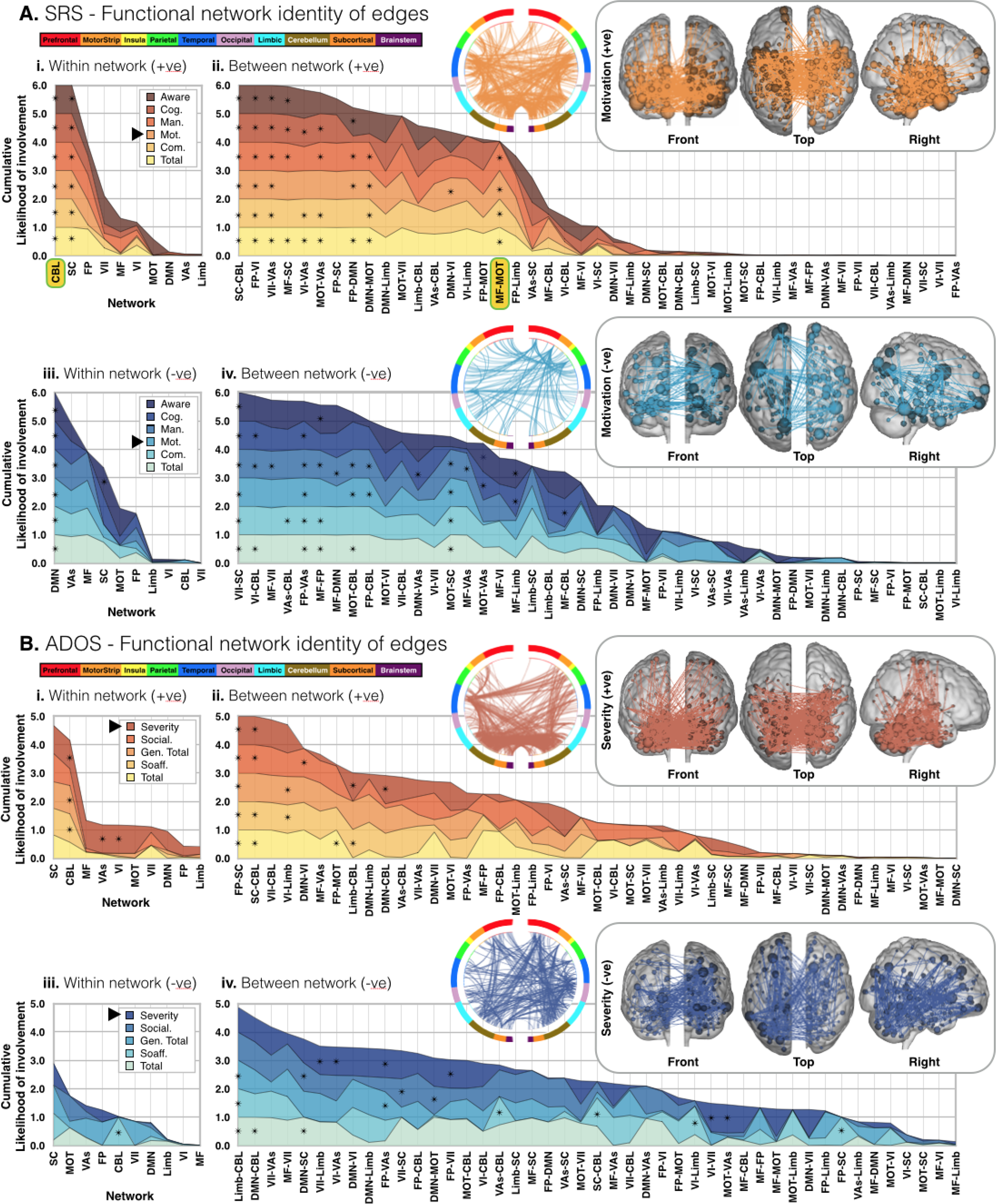
Anatomy of SRS and ADOS sub-scale networks. For SRS (**A.**) and ADOS (**B.**), edge overlap within (**i./iii.**) and between (**ii./iv.**) ten *a priori* atlas networks and our CPM networks are plotted for +ve (**i./ii.**) and -ve (**iii./iv.**) feature sets. Each layered plot shows the cumulative (sum) likelihood (1.0-Pvalue) estimated from the probability of edges being shared between *a priori* networks and each SRS/ADOS sub-scale network. Likelihoods greater than chance are indicated with an asterisk. Notice that in all plots, networks, and inter-network pairs, are ordered from greatest to least cumulative likelihood (i.e. the x-axis is ordered differently in each plot). Inlays show the edges of example SRS/ADOS +ve/-ve sub-scale networks as circle-plots as well as edges/nodes overlaid on glass brains.

### 3.3 Anatomy of SRS/ADOS networks

Unsurprisingly, given that sub-scale scores were highly correlated [Supplementary Figure 1.A.], the anatomy of sub-scale networks was found to be largely similar (e.g. across all SRS sub-scales, +ve edges are very likely to overlap with edges within the cerebellum). On the other hand, there were notable exceptions where sub-scale network anatomy diverged (e.g. edges between medial-frontal and motor networks were very likely to occur within SRS total, communication, motivation and mannerism +ve sub-scale networks, but unlikely to occur within SRS cognition and awareness +ve sub-scale networks) [Figure 3. examples highlighted]. Although it is difficult to summarize the complex networks generated with CPM, here feature sets which contribute most to SRS and ADOS networks are described in a more familiar framework.

### 3.4 Composite SRS/ADOS networks

As a data reduction strategy before investigating model generalizability, and to identify edges that contribute across sub-scales, ’low’ to ’high’ threshold, ’composite’ networks were defined as follows: lowest - edges which appeared in any sub-scale network *at least once*, to highest - edges appear in *all* sub-scale networks. Note that this is not a threshold applied at the feature selection step, but at the level of comparing networks for cross sub-scale relevance. The anatomy of composite networks across thresholds is summarized in Supplementary Figure 5. Despite similar anatomy at the network-level between sub-scale networks, at the edge-level, there was an order of magnitude difference in the number of edges contained within composite networks at the lowest versus highest threshold [Supplementary Figure 6]. However, the anatomy and distribution of edge lengths (ref. below 3.5) in composite networks was similar across thresholds. The feature that distinguished edges in the low-from those in the high-threshold composite networks was the magnitude of the slope in the linear model relating edge strength to clinical score. However, composite network predictive power changed very little with threshold, which is notable considering the difference in the number of edges between thresholds [Supplementary Figure 6.C.]. For all between scale comparisons (ref. below 3.6 and 3.7), composite networks were formed with edges that appeared in at least three sub-scale networks.

### 3.5 Edge lengths

Motivated by controversy in the literature regarding long/short-range hyper/hypo-connectivity in ASD, we analyze the distribution of edge lengths in +/-ve sub-scale and composite networks [Supplementary Figure 6.A./B.]. None of our networks contain outliers. For networks that were not normally distributed, edges skewed towards longer lengths. For both sub-scale and composite SRS networks, there was no difference in median edge length between the +ve and -ve networks. On the other hand, although the difference was small (∼0.5cm), -ve were longer than +ve edge lengths in most sub-scale and composite ADOS networks. In summary, we found weak evidence of longer edges contributing more to symptom severity in ASD.

### 3.6 Model generalizability (ADOS vs SRS)

Model generalizability was tested within the ABIDE-I/II data set, and across data sets and diagnosis (ref. below 3.7). Within ABIDE-I/II, SRS composite networks were applied to individuals from ABIDE-I/II for whom ADOS but not SRS scores were available. Likewise, ADOS composite networks were applied to individuals with SRS but without ADOS scores. Predicted SRS scores correlated with known ADOS social affect (R=0.36, P<0.01), and generic total (R=0.29, P<0.03) scores. Likewise, predicted ADOS scores correlated with known SRS mannerisms (R=0.16, P<0.01) and cognition scores (R=0.20, P<2E-03) [Figure 5.A.].

**Figure 4.**
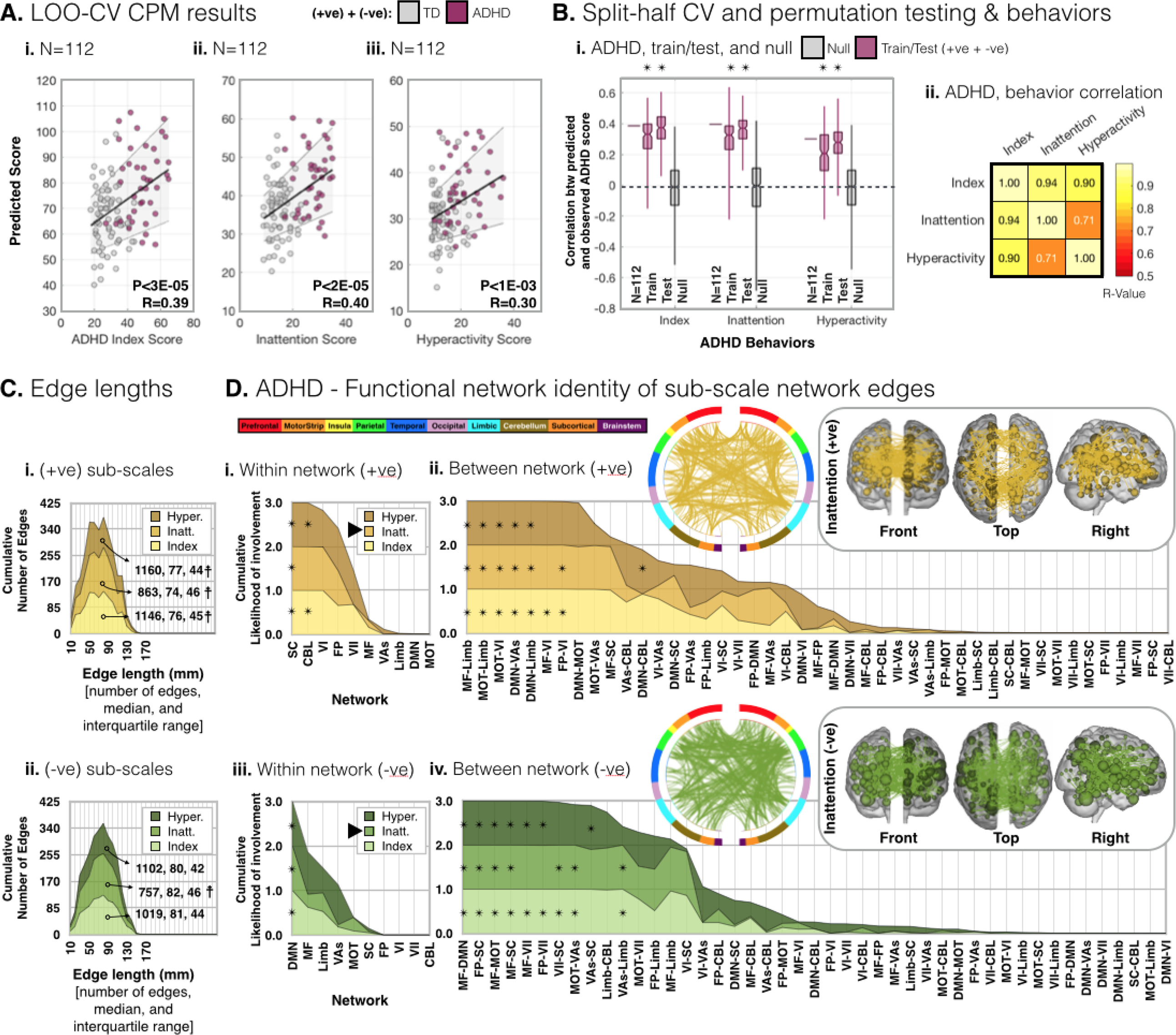
Results from ADHD/TD individuals from the ADHD -200 data set. As in Figure 1.A./B., (**A.**) LOO-CV CPM results for ADHD sub-scale scores. For each sub-scale (**i.-iii.**) the sum of the predicted ADHD score from the +ve/-ve models are plotted against known score. As in Figure 2.A., (**B.i.**) Correlation (R-value) of split-half CV CPMs (n=200) for each ADHD sub-scale and null results from shuffled data (n=1,000). As in Supplementary Figure 1.A., (**B.ii.**) correlation matrix of ADHD behavior sub-scale scores. As with SRS and ADOS, ADHD sub-scale scores are highly correlated. As in Supplementary Figure 6.A.i./B.i., (**C.**) shows layer plots of the cumulative number of edges versus edge length for ADHD sub-scale networks. Networks with edge lengths which are not normally distributed are denoted by a cross (☨). For all not normally distributed networks, edges are skewed towards longer lengths. None of the networks are prone to outliers. There is a difference between +ve and -ve feature set edge lengths for all sub-scale networks (P<4E-03). As in Figure 3., (**D.**) ADHD edge overlap within (**i./iii.**) and between (**ii./iv.**) ten *a priori* atlas networks and ADHD networks are plotted for +ve (**i./ii.**) and -ve (**iii./iv.**) feature sets. Inlays show the edges of example sub-scale networks as circle-plots as well as edges/nodes overlaid on glass brains.

**Figure 5.**
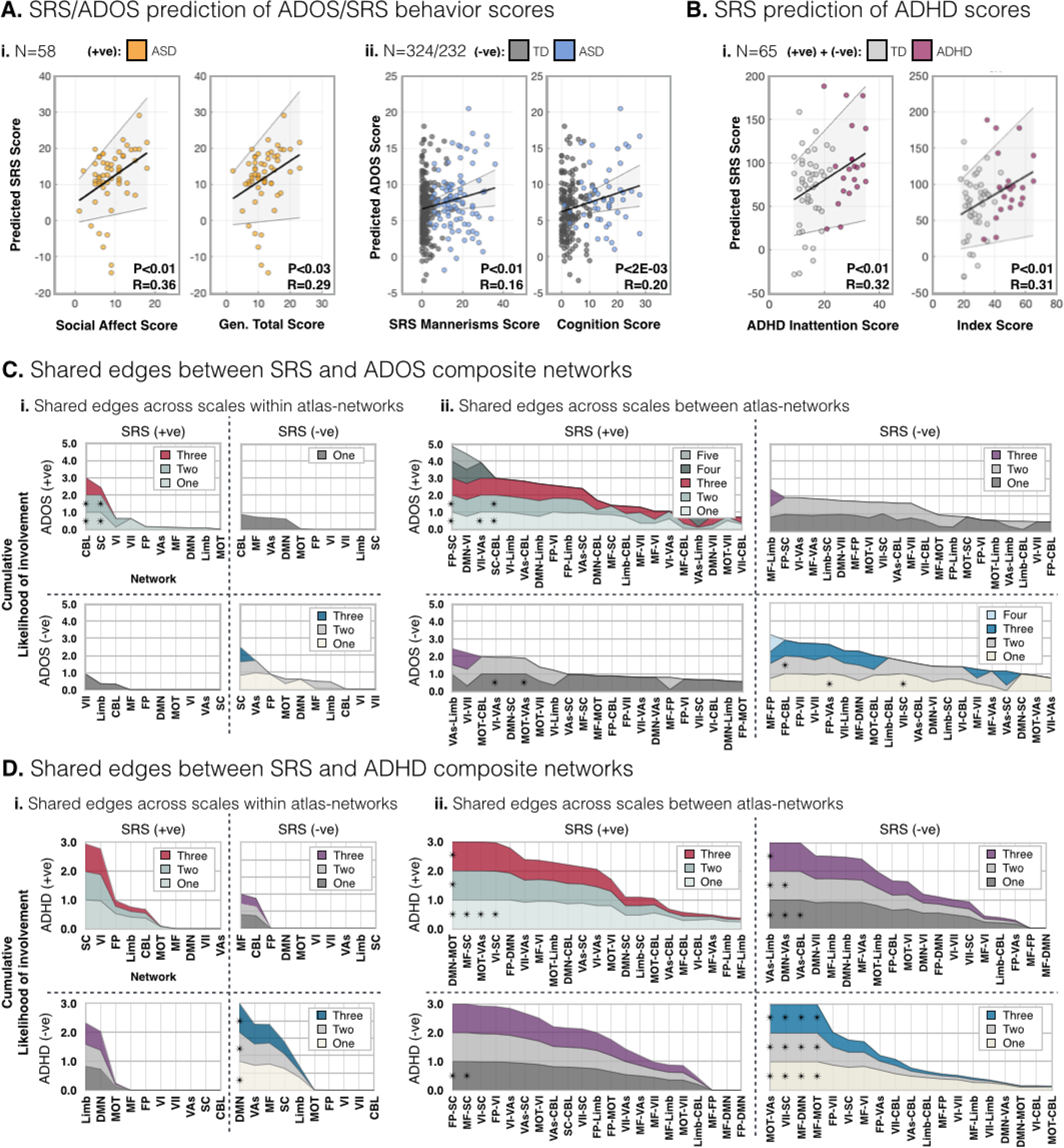
Generalizability of composite networks and overlap of composite network edges: SRS and ADOS (+ve/-ve) & SRS and ADHD (+ve/-ve) Plotted in (**A.**) and (**B.**) are correlations of predicted versus known behavior using composite networks applied across scales. All composite networks were thresholded at three. In (**A.**), the SRS (**i.**) and ADOS (**ii.**) composite networks were used to predict scores for individuals from ABIDE-I/II for whom only the other score was available (i.e. the SRS network was used to predict scores for individuals for whom ADOS scores (not SRS scores) were available). Composite networks were also applied across the ABIDE-I/II and ADHD-200 data sets. (**B.**) Predicted SRS scores correlate with known ADHD scores in individuals from the ADHD-200 data set. Layer plots showing the shared anatomy of SRS and ADOS (**C.**) and SRS and ADHD (**D.**) composite networks across thresholds. Composite network overlap of +ve and -ve feature sets was computed by taking the products: +ve/+ve (upper left, red/grey), +ve/-ve (upper right, purple/grey), -ve/+ve (lower left, purple/grey), and -ve/-ve (lower right, blue/grey) of paired networks and computing the likelihood that each atlas network contribute the observed number of edges to each set of shared features.

### 3.7 Model generalizability (SRS and ADOS models in ADHD)

Motivated by the idea that the underlying biology of mental health disorders is not merely categorical, but rather trans-diagnostic, we tested the SRS and ADOS CPMs to assess their specificity to ASD by applying the models to another neurodevelopmental cohort: children with ADHD.^53–56^ To facilitate this comparison, we first implement the same procedures described above for ABIDE-I/II data, to predict ADHD symptoms within the ADHD-200 data set [Figure 4.A.]. Correlations between known and predicted scores were found to be significant using split-half CV and permutation testing [Figure 4.B.]. As above, we also examined the anatomy of ADHD sub-scale [Figure 4.C.D.] and composite [Supplementary Figure 7.] networks.

Across neurodevelopmental disorders, SRS and ADOS composite networks were applied to individuals from the ADHD-200 data set, and ADHD composite networks applied to ABIDE-I/II data. Predicted SRS scores correlated with known ADHD score (R=0.31/32, P<0.01) [Figure 5.B.]. This result indicates that the SRS model contains components related to attention that accounts for significant variance in predicting ADHD in a different population. However, in each cross-index model test (present and previous section), predictive power was worse than the model constructed with the score of interest.

### 3.8 Shared anatomy of composite networks across scales

To investigate whether predictions across scales were a byproduct of common anatomy, shared features were quantified. The anatomy of shared edges was estimated by taking the products of composite network pairs (SRS/ADOS, and SRS/ADHD) and computing the likelihood that each of the resulting sets of shared features contained *n* edges from atlas-networks. These results are summarized in 2x2 matrices of layer plots for all thresholds [Figure 5.C./D.]. Shared network-level features were summarized and compared to shared edge-level features [Supplementary Figure 6]. For each composite network, the contributing atlas-networks and atlas-network pairs were listed. Common atlas-network features are indicated between composite networks, as are features implicated at the edge-level. Broadly, the cerebellum contributes to SRS and ADOS (+ve), the frontal-parietal to visual areas network pair contributes to SRS and ADOS (-ve), the subcortical network and the frontal-parietal to visual-I network pairs contribute to SRS and ADHD (+ve), and the default mode network contributes to SRS and ADHD (-ve). However, only SRS and ADHD (-ve) share edges that contribute significantly to both networks.

## DISCUSSION

Using a large sample of open-source data and a novel prediction framework, we find meaningful patterns of functional connectivity that can independently predict two clinical measures of ASD symptom severity: ADOS and SRS. In addition, we show that the SRS network predicts symptom severity for another developmental mental health disorder, ADHD. This observation is consistent with a growing body of literature suggesting that ASD and ADHD contain partially overlapping but independent comorbidities, sharing a continuous spectrum of impairment.^53–56^ To ensure that these relationships are not simply the byproduct of a high overlap between networks, we show that <2% of edges are shared across SRS/ADOS/ADHD networks.

In line with previous CPM results, our predictive networks are complex and distributed across the whole brain. Thus, they are not easily described in terms of traditional functional networks. However, some of the same feature sets are implicated across networks and include many areas already identified in the ASD literature: default mode, limbic, visuo-spatial, motor, subcortical, and cerebellum regions.^4, 19–23, 26, 27^ However, with our approach, we cannot conclude that one or a few networks ‘cause’ ASD symptoms but instead observe a convergence of functional connections related to a spectrum of behaviors. We assert that this is indeed a strength of the CPM approach, which affords the ability to resolve more nuanced information by requiring fewer statistical tests. Furthermore, our methodology is intentionally designed to model the underlying biology of mental health disorders as a continuous spectrum not merely a categorical definition. Such models can also be trans-diagnostic, and the results shown here support this hypothesis.^57^

Our study has several limitations. One of which is our strict inclusion criteria. On one hand, we include individuals on medication and both sexes in an attempt to reflect the true patient population, and because we determined clinical scores are independent of medication status and sex. On the other hand, because age, FIQ, and motion are significantly correlated with clinical scores, we are obligated to limit these attributes to eliminate nuisance effects and uncover the connectivity features that relate only to the clinical measures of interest. Another consideration is the inherent heterogeneity of the ABIDE-I/II data set (different sites, acquisition protocols, and behavioral questionnaires and clinical scoring), which we could not account for and likely made prediction more challenging. Nevertheless, that the CPM approach works despite these challenges should be considered a strength. In addition, this heterogeneity may also have strengthened the generalizability. Our models capture ∼10-45% of the variance. In part, this is likely due to differences in sites, acquisition protocols, and behavioral questionnaires. It could also be due to the limited amount of resting-state data available for each subject and noise in the data due to many factors including motion, individual anatomical/functional differences, brain state, a range of physiological variables that can influence connectivity, and the inherent heterogeneity within the patient population. Recent studies have suggested that more data (longer resting-state acquisitions) are needed for high reliability in single subject connectivity assessments.^51, 58^ Finally, it should be noted that the correlative relationships between the functional connectome and clinical scores revealed by CPM cannot be used to infer causality.

Future studies could be improved by implementing longer imaging times and harmonized scanners, to provide more reliable functional connectivity measurements.^51, 58^ It has also been suggested that connectivity data obtained while the subject performs a specific task aimed at enhancing differences in connectivity can lead to better predictive models.^15, 59^ The use of naturalistic conditions such as movie-watching can improve head motion, and tolerance of longer scan durations while enhancing individual differences.^60, 61^ Such conditions may prove particularly advantageous in neurodevelopmental populations as in the current study.

In conclusion, the present work uses a data-driven approach to develop objective quantitative models that establish a link between the individual functional connectome and behavior in ASD. We observe widespread differences in functional organization in individuals with ASD, congruent with the complex behavioral and cognitive abnormalities that are a hallmark of the autism spectrum. We also demonstrate the generalizability and trans-diagnostic utility of this approach. In the future, understanding the changes in functional organization of the brain related to various dimensional aspects of ASD may provide the needed inferential leverage at the individual level to change treatment strategies for ASD individuals and their families.

## Acknowledgements

**ABIDE-I,** primary support for the work by Adriana Di Martino was provided by the NIMH (K23MH087770) and the Leon Levy Foundation. Primary support for the work by Michael P. Milham and the INDI team was provided by gifts from Joseph P. Healy and the Stavros Niarchos Foundation to the Child Mind Institute, as well as by an NIMH award to MPM (R03MH096321). **ABIDE-II,** primary support for the work by Adriana Di Martino and her team was provided by the National Institute of Mental Health (NIMH 5R21MH107045). Primary support for the work by Michael P. Milham and his team provided by the National Institute of Mental Health (NIMH 5R21MH107045); Nathan S. Kline Institute of Psychiatric Research). Additional Support was provided by gifts from Joseph P. Healey, Phyllis Green and Randolph Cowen to the Child Mind Institute. **ADHD-200**, coordinated by Michael P. Milham, M.D., Ph.D. Data collection at Peking University was supported by the following funding sources: The Commonwealth Sciences Foundation, Ministry of Health, China (200802073); The National Foundation, Ministry of Science and Technology, China (2007BAI17B03); The National Natural Sciences Foundation, China (30970802); The Funds for International Cooperation of the National Natural Science Foundation of China (81020108022); The National Natural Science Foundation of China (8100059); Open Research Fund of the State Key Laboratory of Cognitive Neuroscience and Learning.

## Supplementary Material

### Imaging summary

**ABIDE-I/II.** Data was acquired on either a Phillips, GE, or Siemens 3.0T scanner (with the exception of Institut Pasteur, where data was acquired on a Phillips 1.5T scanner). Mean ± standard deviation (SD) and range (min.-max.) of acquisition parameters across institutes are: TR/TE 2083 ± 635 (475-3000) / 29 ± 4 (24-45) msec., number of frames 242 ± 208 (85-947), inplane 3.2 ± 0.4 (2.5-3.8) mm2, through-plane resolution 3.5 ± 0.5 (2.5-4.0) mm, and number of slices 40 ± 6 (31-50).

**ADHD-200.** Data was acquired on a Siemens 3.0T scanner as follows: TR/TE 2000/30 msec., in-plane resolution 2mm2, through-plane resolution 3.0/3.5mm, and number of slices 33. Each 8-minute (240 frame) acquisition was repeated three times during each session. Data were concatenated across acquisitions for connectivity analysis.

**Sup. Figure 1.**
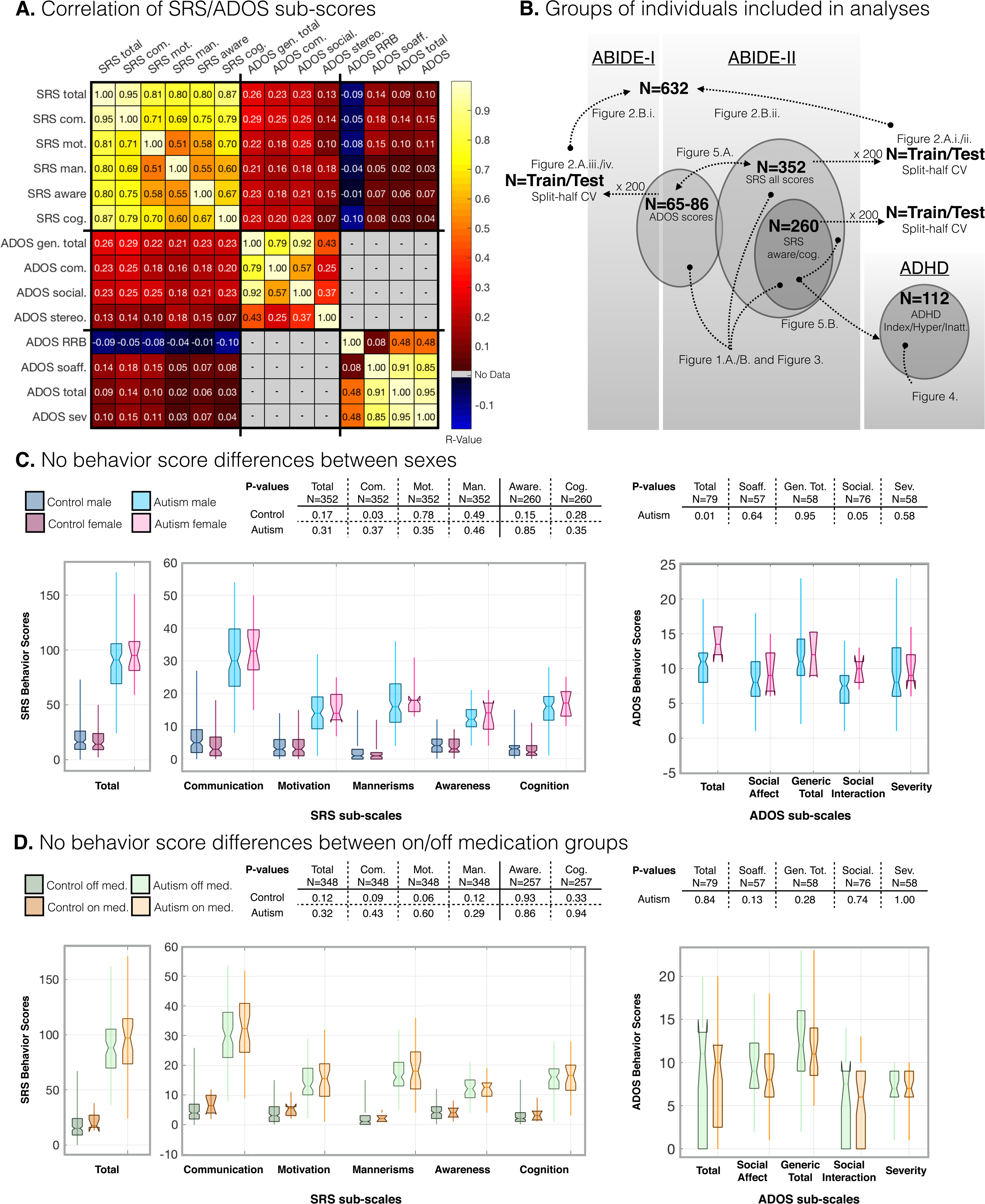
Behavior metrics, data sets & groups, and sex/medication status dependence of SRS/ADOS behavior scores.

**Sup. Figure 2.**
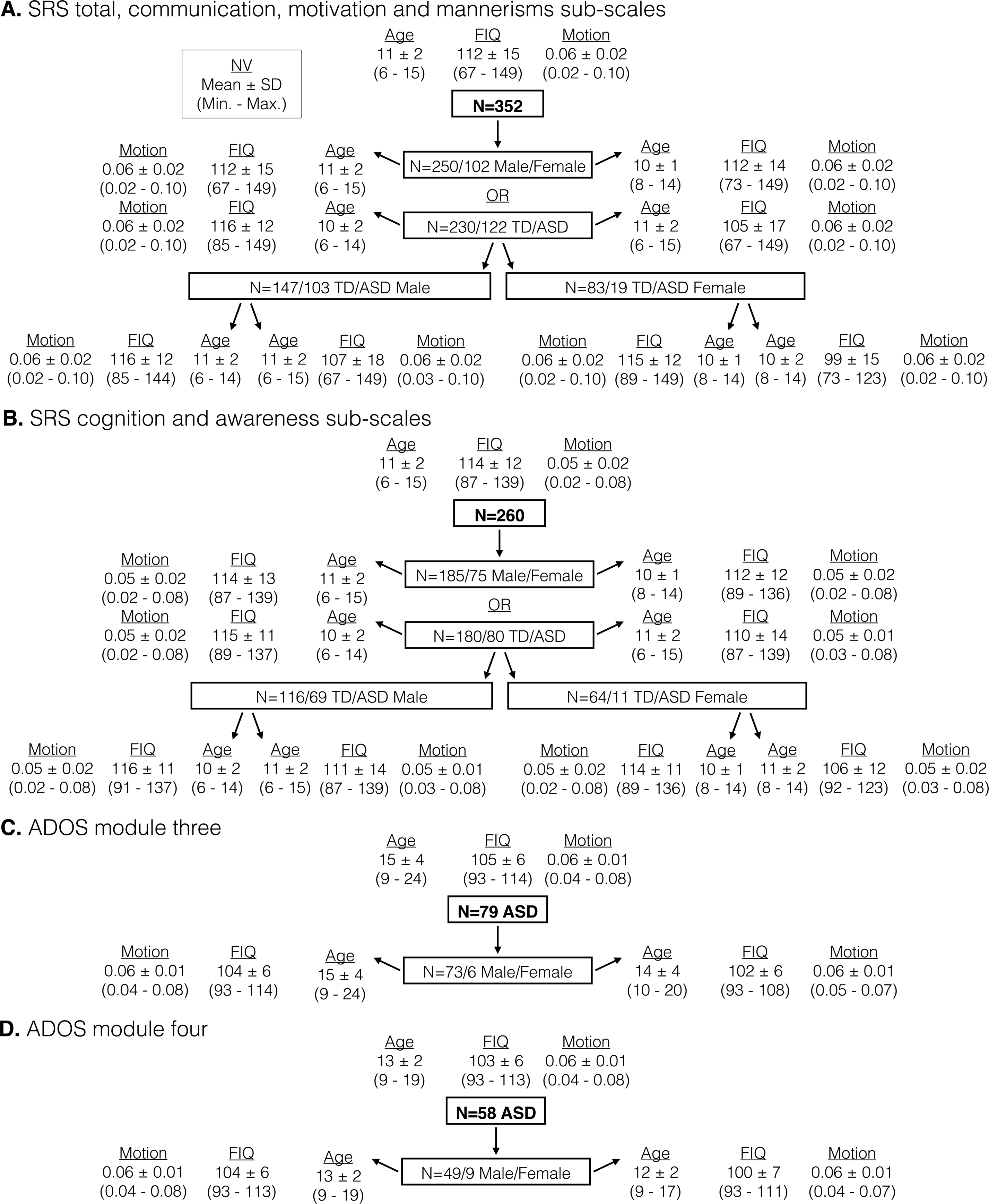
Mean ± SD (min.-max.) motion, FIQ and age for male/female, TD/ASD groups.

**Sup. Figure 3.**
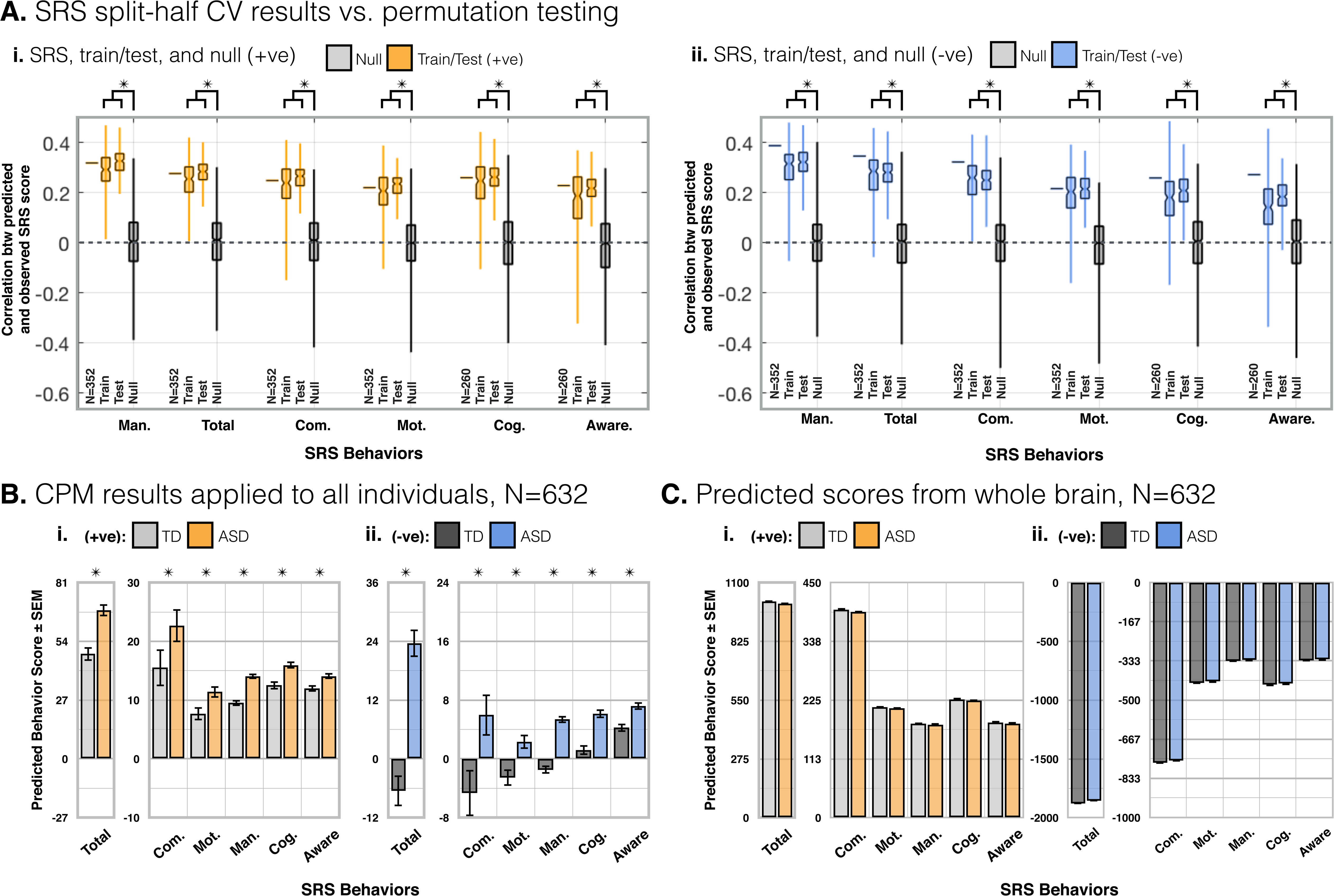
(+ve/-ve) SRS split-half CV, permutation testing results, and whole brain connectivity predicted SRS/ADOS scores in all individuals. **A.** As in Figure 2.A.i. for +ve **(i.)** and -ve **(ii.)** feature sets. **B.** As in Figure 2.B.i. for +ve and -ve feature sets (P<1E-19). **C.** As a control, replacing CPM connectivity measures with whole brain connectivity (all edges) to predict behavior scores for all individuals (N=632) results in no difference between ASD and TD individuals (P>0.3).

**Sup. Figure 4.**
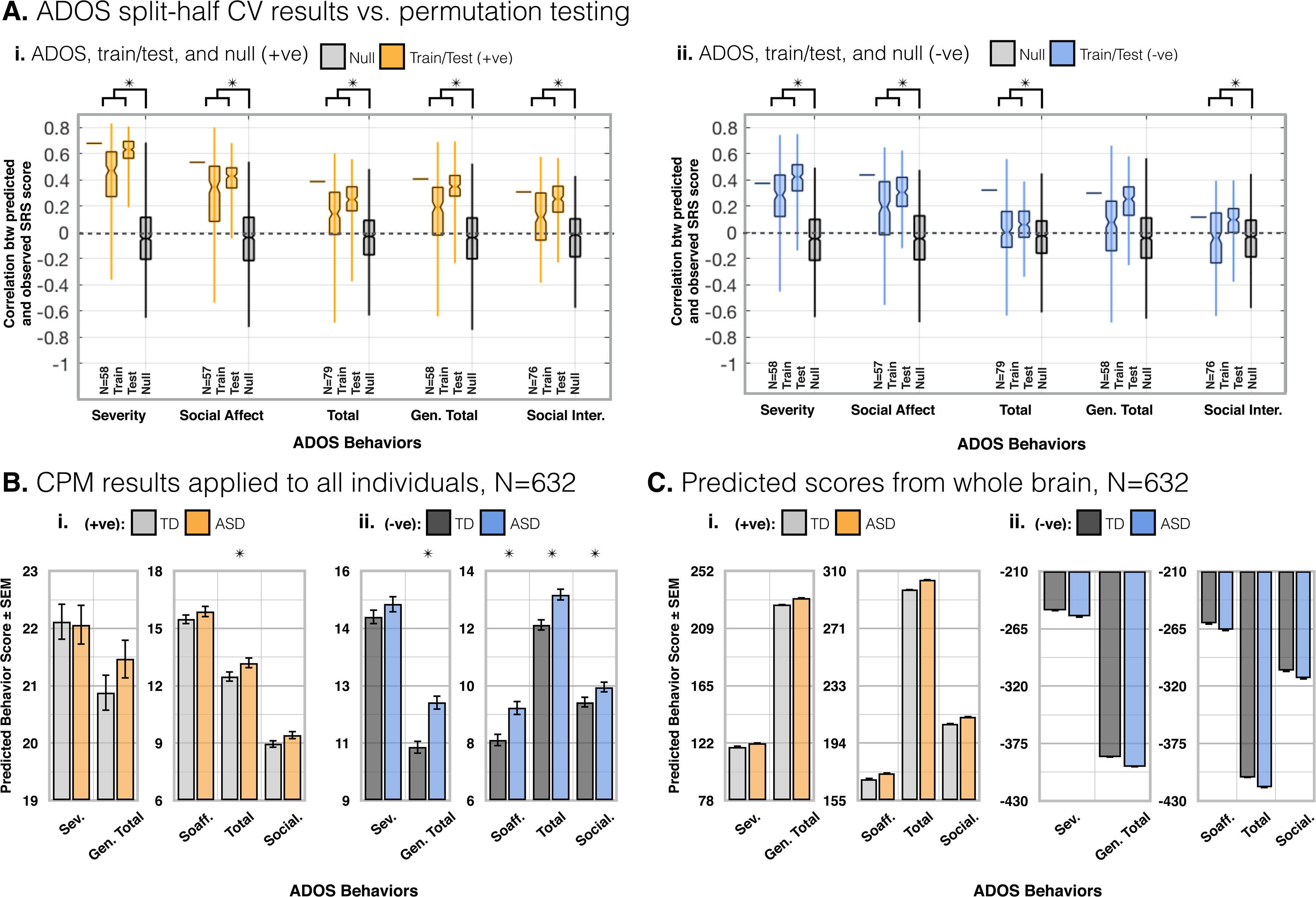
(+ve/-ve) ADOS split-half CV, permutation testing results, and whole brain connectivity predicted ADOS scores in all individuals. **A.** As in Sup. Figure 2.A.ii. +ve **(i.)** and -ve **(ii.)** feature sets. **B.** As in Figure 2.B.ii. for +ve and -ve feature sets (P<0.01). **C.** As a control, replacing CPM connectivity measures with whole brain connectivity (all edges) to predict behaviour scores for all individuals (N=632) results in no difference between ASD and TD individuals (P>0.3).

**Sup. Figure 5.**
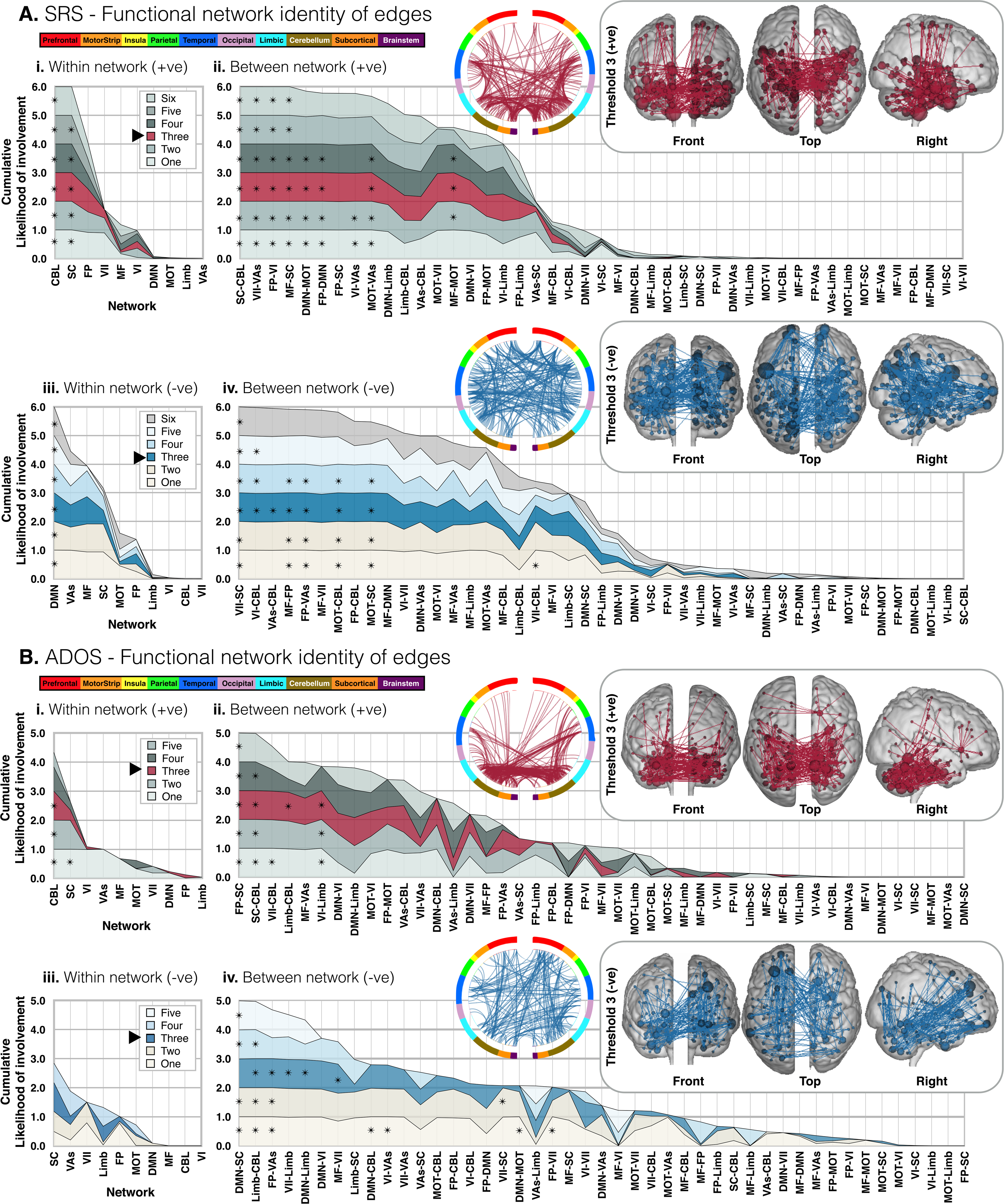
Anatomy of SRS and ADOS composite networks. As in Figure.3., for SRS and ADOS, layered plots show the cumulative likelihood estimated from the probability of edges being shared between *a priori* atlas networks and each SRS or ADOS composite network. Significant overlap (>0.9991) is indicated with asterisks (✴). Here, in place of sub-scale networks [Figure 3.], composite network thresholds are plotted. For SRS composite networks **(A.)**, ‘one’ is the lowest threshold (edges appearing at least once in any sub-scale network, and ‘six’ is the highest threshold (edges appearing in all sub-scale networks). For ADOS **(B.)**, ‘one’ is also the lowest threshold, and ‘three’ is the highest threshold (there being only three ADOS sub-scale networks). Inlays show the edges of example SRS/ADOS +ve/-ve composite networks (threshold = 3) as circle-plots as well as edges/nodes overlaid on glass brains.

**Sup. Figure 6.**
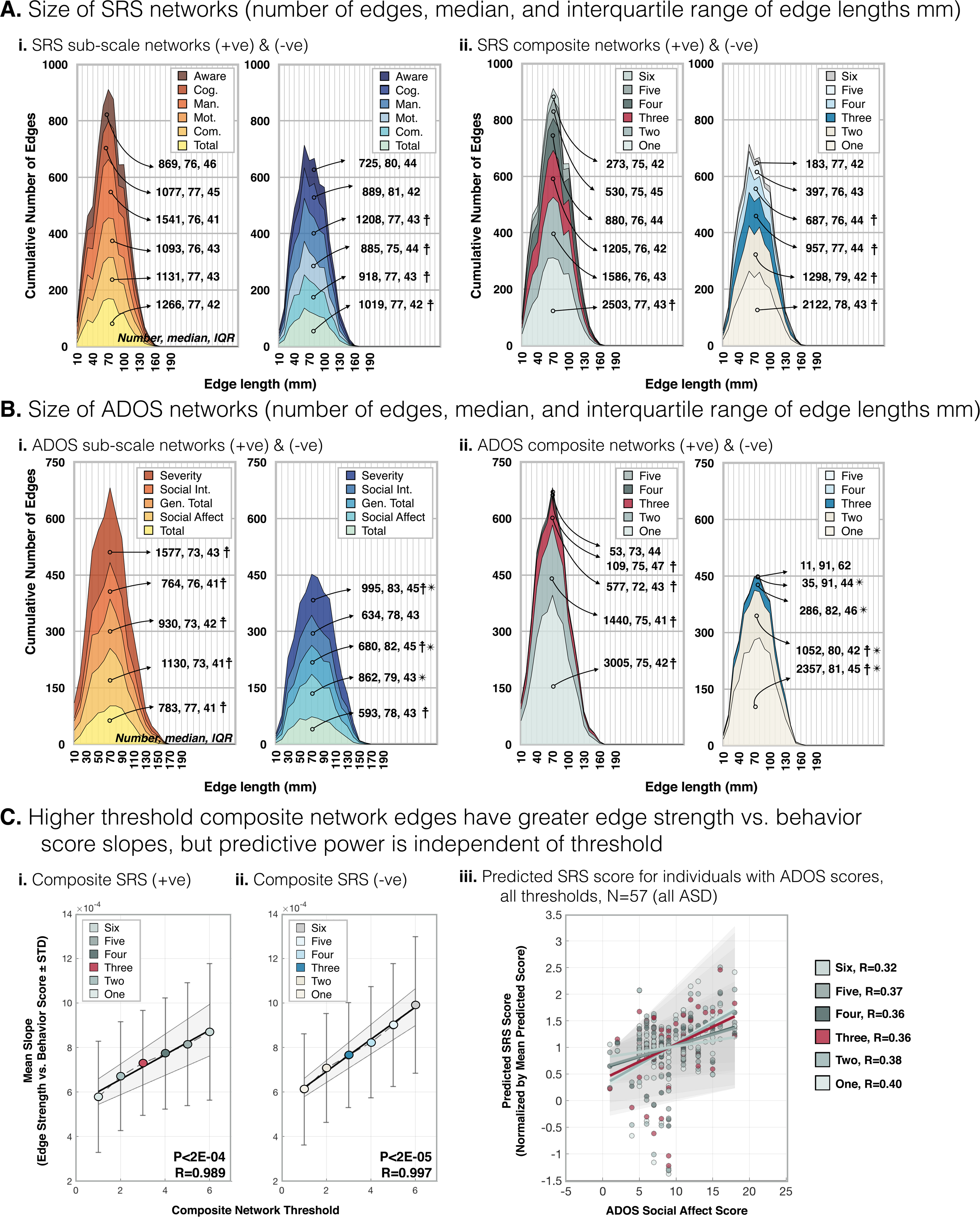
Edge features of SRS/ADOS sub-scale and composite networks. For SRS **(A.)** and ADOS **(B.)** layer plots of the cumulative number of edges in sub-scale **(A./B.i.)** and composite **(A./B.ii.)** networks versus edge length. For each sub-scale and composite network, the number of edges as well as the median and interquartile range of edge lengths (mm) is indicated on each layer plot. Sub-scale networks **(A./B.i.)**, have a similar number of edges. On the other hand, there is a large discrepancy in the number of edges contained in the lowest versus highest threshold composite network (approximately an order of magnitude difference). Networks with edge lengths which are not normally distributed (*lillietest*, MATLAB) are denoted by a cross (☨). For all not normally distributed networks, edges are skewed towards longer lengths (*skewness,* MATLAB). No networks are prone to outliers (*kurtosis*, MATLAB). There is a difference between +ve and -ve feature set edge lengths (✴) for ADOS sub-scale (P<2E-10) and composite networks (*ranksum,* MATLAB, P<0.02). In summary, edge length distributions are largely consistent between sub-scale and composite networks. The feature which does distinguish between edges contained within low versus high threshold composite networks is the magnitude of the slope of edge strength versus behavior score **(C.i./ii.)** (i.e. edge strengths which change less with differences in behavior score are contained within low-threshold composite networks, and edges which change more are contained within high-threshold composite networks). However, predictive power of composite networks is largely independent of threshold **(C.iii.)**. All composite SRS networks were used to predict SRS scores in an independent sample (N=57, ABIDE-I/II individuals with ADOS, but without SRS scores available). Predicted SRS versus known ADOS score for thresholds one through six are plotted.

**Sup. Figure 7.**
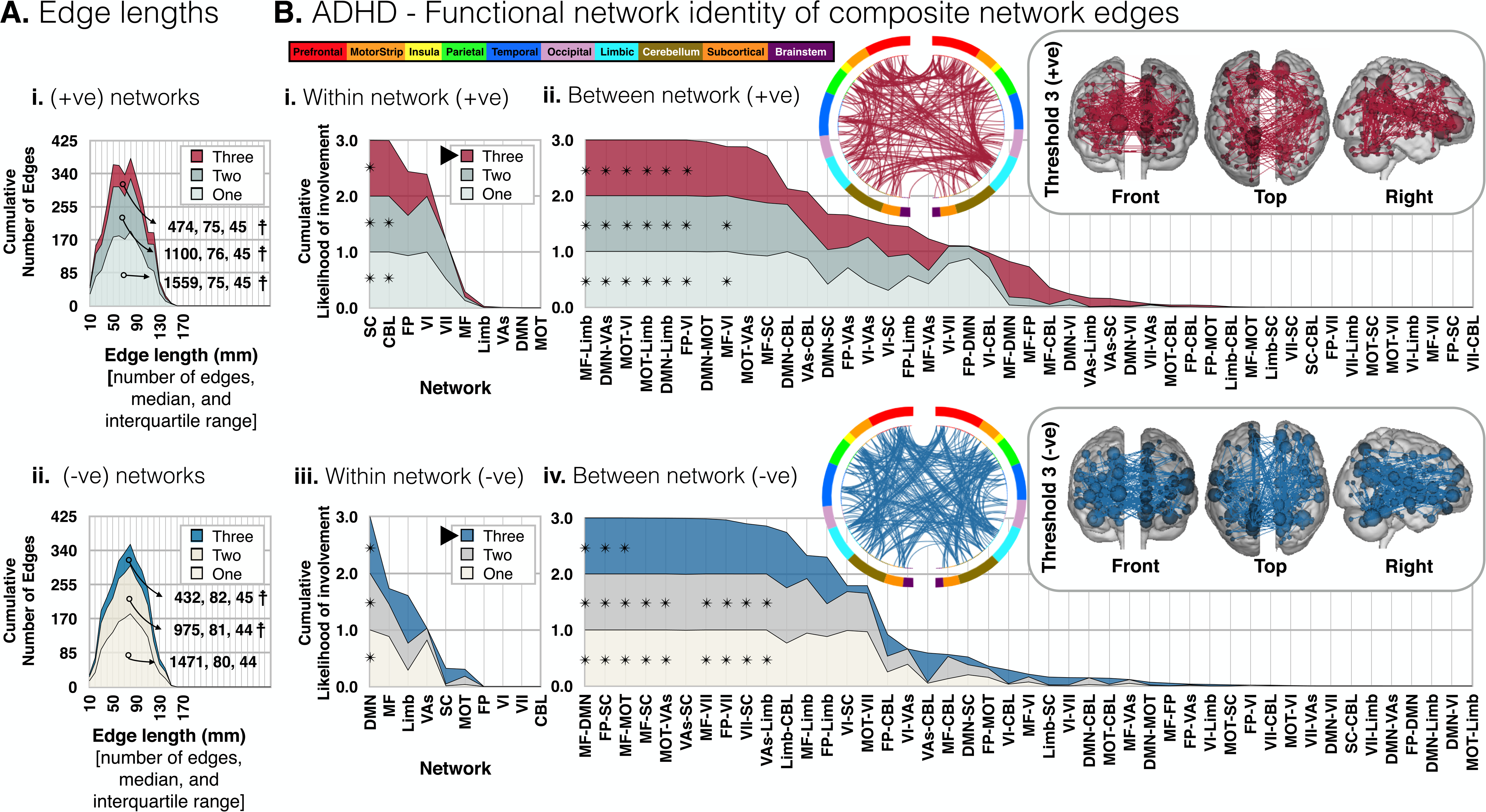
Anatomy of ADHD sub-scale and composite networks. As in Supplementary Figure 4.A.ii./B.ii., **(A.)** shows layer plots of the cumulative number of edges versus length for ADHD composite networks. The number of edges, median and interquartile ranges of edge lengths (mm) is indicated on each layer plot. As with SRS and ADOS composite networks, the number of edges depends on threshold. More edges belong to the lowest threshold and fewer to the higher threshold networks. Networks with edge lengths which are not normally distributed (*lillietest*, MATLAB) are denoted by a cross (☨). For all not normally distributed networks, edges are skewed towards longer lengths (*skewness,* MATLAB). None of the networks are prone to outliers (*kurtosis,* MATLAB), and there is a difference between +ve and -ve feature set edge lengths for all networks (*ranksum,* MATLAB, P<1E-03). As in Supplementary Figure 3., **(B.)** shows layered plots of the cumulative likelihood estimated from the probability of edges being shared between *a priori* networks and each ADHD composite network. Inlays show the edges of example +ve/-ve composite networks (threshold = 3) as circle-plots as well as edges/nodes overlaid on glass brains.

**Sup. Figure 8.**
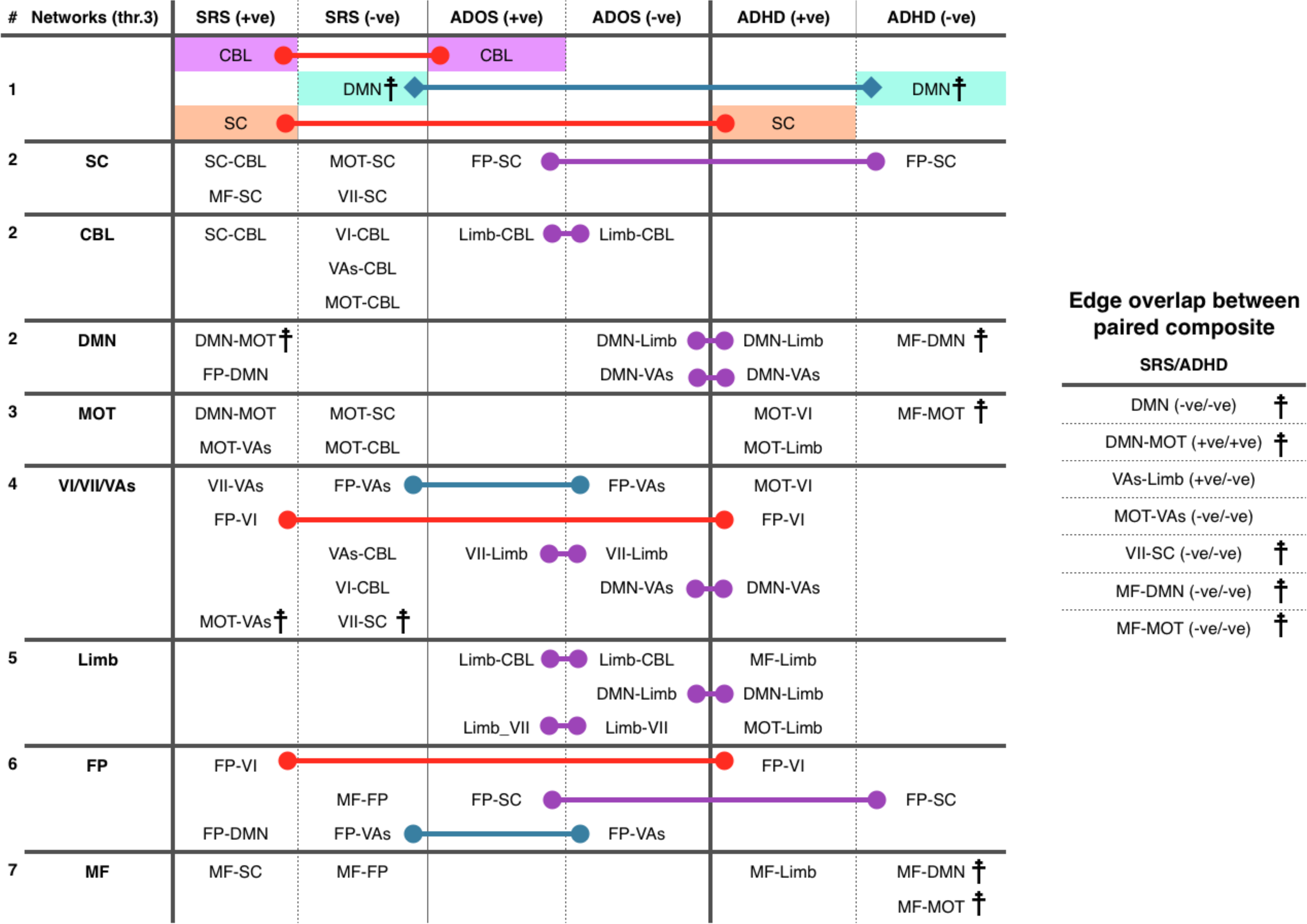
Summary of SRS/ADOS/ADHD composite network anatomy and shared across scale anatomy at the network and edge level. From layer plots in Supplementary Figures 5 (SRS/ADOS) and 7 (ADHD) we infer which are the shared contributing functional network feature sets between composite SRS, ADOS and ADHD networks. We relate this ‘network-level’ summary to the strict edge overlap across scales [Figure 5 C./D.]. We compare at composite network threshold three. In each column in the main table (left), the contributing atlas-networks and atlas-network pairs (feature sets) are listed for each composite network. Contributing feature sets are indicated in the aforementioned Supplementary Figures by an asterisk (✴). In row #1, contributing within-network feature sets are listed: CBL, SC, and DMN. In rows labeled #2, between-network feature sets are listed which include the CBL, SC, and DMN (row #1 networks). Rows labeled #3-7 list all other between-network feature sets. Note that entries are repeated (e.g. SC-CBL is listed twice: once in the ‘SC’, and once in the ‘CBL’ row). Contributing feature sets found in two composite networks are connected by colored lines. If contributing feature sets are found between two +ve composite networks the connecting line is red, if between two -ve composite networks the line is blue, and if between +ve and -ve composite networks the line is purple. Given that a higher score indicates a worse prognosis on all scales, shared features between +ve and -ve networks (purple lines) implies duplicitous roles of these feature sets (i.e. increased network strength is correlated with a worse score on one scale, but a better score on another scale). Recall that not all edges within a contributing feature set are found within any composite network (just more than would be expected by chance). Thus, it does not follow from this result that the same edges are implicated in these instances. All shared features between +ve and -ve composite networks are within ADOS composite networks, or between ADOS and ADHD scales. In our across scale predictions [Figure 5.A.], the ADOS composite network did not predict ADHD scores. Likewise, the ADHD composite network did not predict ADOS scores. Evidence of seemingly opposite contributions by the same underlying feature sets may be in-line with this result. Feature sets identified by the strict edge-overlap between composite networks [Figure 5 C./D.] are listed in the auxiliary table (right). There are no contributing shared edges between SRS and ADOS composite networks. Shared SRS/ADHD edges which also appear in the main table are indicated by a cross (☨). The only feature set identified at both the network and edge level (i.e. shares edges across SRS and ADHD scales and contributes significantly to both composite networks) is the DMN in SRS/ADOS (-ve/-ve), indicated by a square ended line. Although significant, the number of shared edges between SRS/ADHD (-ve/-ve) within the DMN is still very small: <2% of edges within either network.

**Sup. Table 1.** SRS/ADOS (ABIDE-I/II) and ADHD (ADHD-200) CPM results.

A. SRS, CPM results - with age, FIQ and motion as covariates (ref. Figure 1. A.), N=352/260
| SRS<br>(+ve) + (-ve) | Model |  | Age |  | FIQ |  | Motion |  |
| --- | --- | --- | --- | --- | --- | --- | --- | --- |
|  | R-value | P-Value | R-value | P-Value | R-value | P-Value | R-value | P-Value |
| Total Score | 0.32 | 5E-10 | -0.01 | 0.86 | -0.01 | 0.90 | -0.04 | 0.50 |
| Communication | 0.30 | 1E-08 | -0.03 | 0.61 | 0.01 | 0.90 | -0.04 | 0.50 |
| Motivation | 0.23 | 2E-05 | -0.04 | 0.49 | 0.01 | 0.88 | -0.04 | 0.45 |
| Mannerisms | 0.37 | 1E-12 | 0.01 | 0.81 | 0.01 | 0.88 | -0.04 | 0.48 |
| Cognition | 0.27 | 1E-05 | -0.00 | 0.97 | 0.02 | 0.76 | -0.08 | 0.19 |
| Awareness | 0.27 | 1E-05 | 0.01 | 0.84 | -0.00 | 0.98 | -0.05 | 0.46 |

B. ADOS, CPM results (ref. Figure 1. B.), N=58-79
| ADOS | Model |  | FIQ |  | Age |  | Motion |  |
| --- | --- | --- | --- | --- | --- | --- | --- | --- |
|  | R-value | P-Value | R-value | P-Value | R-value | P-Value | R-value | P-Value |
| Generic Module 4 |  |  |  |  |  |  |  |  |
| Total (+ve) + (-ve) | 0.40 | 2E-03 | -0.16 | 0.24 | 0.01 | 0.97 | -0.15 | 0.27 |
| Social Interaction (+ve) | 0.31 | 8E-03 | 0.16 | 0.17 | -0.14 | 0.24 | 0.09 | 0.47 |
| Module 3 |  |  |  |  |  |  |  |  |
| Total (+ve) + (-ve) | 0.43 | 1E-04 | 0.16 | 0.16 | -0.07 | 0.55 | 0.17 | 0.15 |
| Social Affect (+ve) + (-ve) | 0.54 | 3E-05 | -0.13 | 0.35 | 0.14 | 0.30 | -0.05 | 0.74 |
| Severity (+ve) + (-ve) | 0.60 | 1E-06 | -0.01 | 0.93 | 0.12 | 0.38 | -0.13 | 0.35 |

C. SRS, CPM results for male individuals, N=250/185
| SRS<br>(+ve) + (-ve) | Model |  | Age |  | FIQ |  | Motion |  |
| --- | --- | --- | --- | --- | --- | --- | --- | --- |
|  | R-value | P-Value | R-value | P-Value | R-value | P-Value | R-value | P-Value |
| Total Score | 0.25 | 9E-05 | -0.01 | 0.91 | -0.01 | 0.83 | -0.07 | 0.29 |
| Communication | 0.25 | 8E-05 | -0.02 | 0.77 | -0.01 | 0.93 | -0.08 | 0.21 |
| Motivation | 0.18 | 5E-03 | -0.04 | 0.48 | 0.01 | 0.93 | -0.08 | 0.22 |
| Mannerisms | 0.32 | 3E-07 | -0.03 | 0.61 | -0.02 | 0.76 | -0.05 | 0.47 |
| Cognition (+ve) | 0.19 | 0.01 | -0.04 | 0.56 | 0.03 | 0.72 | -0.11 | 0.15 |
| Awareness (+ve) | 0.17 | 0.02 | 0.00 | 0.99 | -0.03 | 0.65 | -0.02 | 0.81 |

D. ADOS, results for male individuals, N=47-73
| ADOS | Model |  | FIQ |  | Age |  | Motion |  |
| --- | --- | --- | --- | --- | --- | --- | --- | --- |
|  | R-value | P-Value | R-value | P-Value | R-value | P-Value | R-value | P-Value |
| Generic Module 4 |  |  |  |  |  |  |  |  |
| Total (+ve) + (-ve) | 0.39 | 7E-03 | -0.23 | 0.13 | 0.05 | 0.74 | -0.13 | 0.39 |
| Module 3 |  |  |  |  |  |  |  |  |
| Total (+ve) + (-ve) | 0.23 | 0.05 | 0.14 | 0.25 | -0.11 | 0.36 | 0.10 | 0.41 |
| Social Affect (+ve) + (-ve) | 0.58 | 3E-05 | -0.21 | 0.17 | 0.17 | 0.26 | -0.11 | 0.48 |
| Severity (+ve) + (-ve) | 0.60 | 1E-05 | -0.12 | 0.44 | 0.13 | 0.39 | -0.19 | 0.20 |

E. ADHD, results for all individuals (ref. Figure 4.A.), N=112
| ADHD<br>(+ve) + (-ve) | Model |  | Age |  | FIQ |  | Motion |  |
| --- | --- | --- | --- | --- | --- | --- | --- | --- |
|  | R-value | P-Value | R-value | P-Value | R-value | P-Value | R-value | P-Value |
| Index | 0.39 | 3E-05 | -0.06 | 0.51 | 0.05 | 0.64 | 0.17 | 0.07 |
| Hyperactivity | 0.30 | 1E-03 | -0.13 | 0.16 | -0.01 | 0.94 | 0.13 | 0.17 |
| Inattention | 0.40 | 2E-05 | -0.04 | 0.70 | 0.05 | 0.61 | 0.17 | 0.08 |

## REFERENCES

1. Baxter AJ, Brugha TS, Erskine HE, Scheurer RW, Vos T, Scott JG. The epidemiology and global burden of autism spectrum disorders. Psychol Med 2015; 45(3): 601–613.

2. Erksine HE, Ferrari AJ, Polanczyk GV, Moffitt TE, Murray CJ, Vos T et al. The global burden of conduct disorder and attention-deficit/hyperactivity disorder in 2010. J Child Psychol Psychiatry 2014; 55(4): 328–336.

3. de Bildt A, Sytema S, Ketelaars C, Kraijer D, Mulder E, Volkmar F et al. Interrelationship between autism diagnostic observation schedule-generic (ADOS-G), autism diagnostic interview-revised (ADI-R0, and the diagnostic and statistical manual of metal disorders (DSM-IV-TR) classification in children and adolescents with mental retardation. J Autism Dev Disord 2004; 34(2): 129–137.

4. Minshew NJ, Williams DL. The new neurobiology of autism: cortex, connectivity, and neuronal organization. Archives of neurology 2007; 64(7): 945–950.

5. Plitt M, Barnes KA, Wallace GL, Kenworthy L, Martin A. Resting-state functional connectivity predicts longitudinal change in autistic traits and adaptive functioning in autism. Proc Natl Acad Sci USA 2015; 112(48):E6699–706.

6. Yahata N, Morimoto J, Hashimoto, Lisi, Shibata, Kawakuno H et al. A small number of abnormal brain connections predicts adult autism spectrum disorder. Nat Commun 2016; 7:7:11254.

7. Emerson RW, Adams C, Nishino T, Hazlett HC, Wolff JJ, Zwaigendaum L, et al. Functional neuroimaging of high-risk 6-month-old infants predicts a diagnosis of autism at 24 months of age 2017; 9(393):1–8.

8. Ogawa S, Lee TM, Kay AR, Tank DW. Brain magneti resonance imaging with contrast dependent on blood oxygenation. Proc Natl Acad Sci USA 1990; 87(24):9868–72.

9. Biswal B, Yetkin FZ, Heughton VM, Hyde JS. Functional connectivity in the motor cortex of resting human brain using echo-planar MRI. Magn Reson Med 1995; 34(4):537–41.

10. Biswal BB, Mennes M, Zuo XN, Gohel S, Kelly C, Smith SM et al. Toward discovery science of human brain function. Proc Natl Acad Sci USA. 2010;107(10):4734–9.

11. Finn ES, Shen X, Scheinost D, Rosenberg MD, Huang J, Cun MM et al. Functional connectome fingerprinting: identifying individuals using patterns of brain connectivity. Nat Neurosci 2015; 18(11): 1664–1671.

12. Rosenberg MD, Finn ES, Scheinost D, Papademetris X, Shen X, Constable RT, Chun MM. A neuromarker of sustained attention from whole-brain functional connectivity. Nat Neurosci 2016; 19(1):165–7.

13. Rosenberg MD, Hsu WT, Scheinost D, Todd Constable R, Chun MM. Connectome-based models predict separable components of attention in novel individuals. J Cogn Neurosci. 2018; 30(2):160–173.

14. Beaty RE, Kenett YN, Christensen AP, Rosenberg MD, Benedek M, Chen Q et al. Robust prediction of individual creative ability from brain functional connectivity. Proc Natl Acad SCi USA 2018; 115(5):1087–1092.

15. Rosenberg MD, Finn ES, Scheinost D, Constable RT, Chunn MM. Characterizing attention with predictive network models. Trends Cogn Sci 2017; 21(4): 290–302.

16. Herane LJ, Mattingley JB, Cocchi L. Functional brain networks related to individual differences in human intelligence at rest. Sci Rep 2016; 6:32328.

17. Uddin LQ, Supekar K, Menon V. Reconceptualizing functional brain connectivity in autism from a developmental perspective. Front Hum Neurosci 2013; 7:458.

18. Lee JM, Kyeong S, Kim E, Cheon KA. Abnormalities of inter- and intra-hemispheric functional connectivity in autism spectrum disorder: a study using autism brain imaging data exchange database. Front Neurosci 2016; 10(191):1–11.

19. Redcay E, Moran JM, Mavros PL, Tager-Flusberg H, Gabrieli JD, Whitfield-Gabrieli S. Intrinsic functional network organization in high-functioning adolescents with autism spectrum disorder. Front Hum Neurosci 2013; 7(573): 1–11.

20. Assaf M, Jagannathan K, Calhoun VD, Miller L, Stevens MC, Sahl R et al. Abnormal functional connectivity of default mode sub-networks in autism spectrum disorder patients. NeuroImage 2010; 53(1): 247–256.

21. Ypma RJ, Moseley RL, Holt RJ, Rughooputh N, Floris DL, Chura LR et al. Default Mode hypoconnectivity underlies a sex-related autism spectrum. Biol Psychiatry Cogn Neurosci Neuroimaging 2016; 1(4): 364–371.

22. Chen CP, Keown CL, Jahedi A, Nair A, Pflieger ME, Bailey BA et al. Diagnostic classification of intrinsic functional connectivity highlights somatosensory, default mode, and visual regions in autism. Neuroimage Clin 2015; 9(8): 238–245.

23. Nielsen JA, Zielinski BA, Fletcher PT, Alexander AL, Lange N, Bigler ED et al. Abnormal lateralization of functional connectivity between language and default mode regions in autism. Mol Autism 2014; 5(8):1–11.

24. Farrant K, Uddin LQ. Atypical developmental of dorsal and ventral attention networks in autism. Dev Sci 2016; 19(4): 550–563.

25. Keehn B, Müller RA, Townsend J. Atypical attentional networks and the emergence of autism. Neurosci Biobehav Rev. 2013; 37(2): 164–183.

26. Cerlinani L, Mennes M, Thomas RM, Di Martino A, Thioux M, Keysers C. Increased functional connectivity between subcortical and cortical resting-state networks in autism spectrum disorder. JAMA Psychiatry 2015; 72(8): 767–777.

27. Gotts SJ, Simmons WK, Milbury LA, Wallace GL, Cox RW, Martin A. Fractionation of social brain circuits in autism spectrum disorder. Brain 2012; 135(Pt 9): 2711–2725.

28. von dem Hagen EA, Stoyanova RS, Baron-Cohen S, Calder AJ. Reduced functional connectivity within and between ‘social’ resting state networks in autism spectrum conditions. Soc Cogn Affect Neurosci 2013; 8(6): 694–701.

29. Nielsen JA, Zielinski BA, Fletcher PT, Alexander AL, Lange N, Bigler ED et al. Multisite functional connectivity MRI classification of autism: ABIDE results. Frontiers in Human Neuroscience. 2013; 7(599): 1–12.

30. Lewis DJ, Evans AC, Pruett Jr. JR, Botteron KN, McKinstry RC, Zwaigenbaum L et al. The emergence of network inefficiencies in infants with autism spectrum disorder. Biological Psychiatry 2017; 82: 176–185.

31. Rane P, Cochran D, Hodge SM, Haselgrove C, Kennedy DN, Frazier JA. Connectivity in autism: a review of MRI connectivity studies. Harv Rev Psychiatry 2015; 23(4): 223–244.

32. Yarkoni T, Westfall J. Choosing prediction over explanation in psychology: Lessons from machine learning. Perspect Psychol Sci 2017; 12(6):1100–1122.

33. Rausch A, Zhang W, Beckmann CF, Buitelaar JK, Groen WB, Haak KV. Connectivity-based parcellation of the amygdala predicts social skills in adolescents with autism spectrum disorder. J Autism Dev Disord 2018; 48(2):572–582.

34. Bernhardt BC, Di Martino A, Valk SL, Wallace GL. Neuroimaging-based phenotyping of the autism spectrum. Curr Top Behav Neurosci 2017;30: 341–355.

35. Hull JV, Jacokes ZJ, Torgerson CM, Irimia A, Van Horn JD. Resting-state functional connectivity in autism spectrum disorder: A review. Front Psychiatry 2017; 7:205.

36. Nunes AS, Peatfield N, Vakorin V, Doesburg SM. Idiosyncratic organization of cortical networks in autism spectrum disorder. Neuroimage 2018; S1053-8119(18)30022-3.

37. Shen X, Finn ES, Scheinost D, Rosenberg MD, Chun MM, Papademetris X et al. Using connectome-based predictive modeling to predict individual behavior from brain connectivity. Nat Protoc 2017; 12(3):506–518.

38. Di Martino A, Yan CG, Li Q, Denio E, Castellanos FX, Alaerts K et al. The autism brain imaging data exchange: towards a large-scale evaluation of the intrinsic brain architecture in autism. Mol Psychiatry 2014; 19(6):659–67.

39. Di Martino A, O’Connor D, Chen B, Alaerts K, Anderson JS, Assaf M et al. Enhancing studies of the connectomre in autism using the aytusm imaging data exchange II. Sci Data 2017; 4:170010.

40. Consortium T A.-200. The ADHD-200 Consortium. A Model to Advance the Translational Potential of Neuroimaging in Clinical Neuroscience. Front Syst Neurosci 2012;6:62.

41. Insel T, Cuthbert B, Garvey M, Heinssen R, Pine DS, Quinn K, Sanislow C, Wang P. Research domain criteria (RDoC): toward a new classification framework for research on mental disorders. Am J Psychiatry 2010; 167(7):748–51.

42. Scheinost D, Papademetris X, Constable RT. The impact of image smoothness on intrinsic functional connectivity and head motion confounds. Neuroimage 2014;95:13–21.

43. Joshi A, Scheinost D, Okuda H, Belhachemi D, Murphy I, Staib LH et al. Unified framework for development, deployment and robust testing of neuroimageing algorithms. Neuroinformatics 2011;9(1):69–84.

44. Shen X, Tokoglu F, Papademetris X, Constable RT. Groupwise whole-brain parcellation from resting-state fMRI data for network node identification. Neuroimage 2013 82:403–15.

45. Constantino JN, Pryzbeck T, Friesen D, Todd RD (2000), Reciprocal social behavior in children with and without pervasive developmental disorders. J Dev Behav Pediatr 2000;21(1):2–11.

46. Constantino JN, Todd RD. Autistic traits in the general population: a twin study. Arch Gen Psychiatry 2003;60(5):524–530.

47. Constantino JN, Davis SA, Todd RD, Schindler MK, Gross MM, Brophy SL et al. Validation of a brief quantitative measure of autistic traits: comparison of the social responsiveness scale with the Autism Diagnostic Interview-Revised. J Autism Dev Disord 2003;33(4):427–33.

48. Lord C, Risi S, Lambrecht L, Cook EH Jr, Leventhal BL, DiLavore PC, et al. The autism diagnostic observation schedule-generic: a standard measure of social and communication deficits associated with the spectrum of autism. 2000;30(3):205–23.

49. DuPaul GJ, Power TJ, Anastopoulos AD, Reid R. ADHD Rating Scale-IV: Checklists, norms, and clinical interpretation. Guilford Press; New York 1998:25.

50. Rubinov M, Sporns O. Complex network measures of brain connectivity: uses and interpretations. Neuroimage 2010;52:1059–1069.

51. Noble S, Spann MN, Tokoglu F, Shen X, Constable RT, Scheinost D. Influences of the test-retest reliability of functional connectivity MRI and its relationship with behavioral utility. Cereb Cortex 2017;27(11):5415–5429.

52. Power JD, Barnes KA, Snyder AZ, Schlaggar BL, Petersen SE. Spurious but systematic correlations in functional connectivity MRI networks arise from subject motion. Neuroimage 2012;59(3):2142–54.

53. Taurines R, Schwenck C, Westerwald E, Sachse M, Siniatchkin M, Freitag C. ADHD and autism: differential diagnosis or overlapping traits? A selective review. Atten Defic Hyperact Disord 2012;4(3):115–39.

54. Craig F, Margari F, Legrottaglie AR, Palumbi R, de Giambattista C, Margari L. A review of executive function deficits in autism spectrum disorder and attention-deficit.hyperactivity disorder. Neuropsychiatr Dis Treat 2016;12:1191–202.

55. Proal E, González-Olvera J, Blancas ÁS, Chalita PJ, Castellanos FX. Neurobiology of autism and attention deficit hyperactivity disorder by means of neuroimaging techniques: convergences and divergences. Rev Neurol 2013;6:57.

56. Grzadzinski R, Di Martino A, Brady E, Mairena MA, O’Neale M, Petkova E, et al. Examining autistic traits in children with ADHD: does the autism spectrum extend to ADHD? J Autism Cev Disord 2011; 41(9):1178–91.

57. Insel TR. The NIMH research domain criteria (RDoC) project precision medicine for psychiatry. Am J Psychiatry 2014;171(4):395–7.

58. Noble S, Scheinost D, Finn ES, Shen X, Papademetris X, McEwen SC et al. Multisite reliability of MR-based functional connectivity. Neuroimage 2017;146:959–970.

59. Finn ES, Scheinost D, Finn DM, Shen X, Papademetris X, Constable RT. Can brain state be manipulated to emphasize individual differences in functional connectivity. Neuroimage 2017;160:140–151.

60. Vanderwal T, Kelly C, Eibott J, Mayes LC, Castellanos FX. Inscapes: a movie paradigm to improve compliance in functional magnetic resonance imaging. Neuroimage 2015;122:222–32.

61. Vanderwal T, Eilbott J, Finn ES, Craddock RC, Turnbull A, Castellanos FX. Individual difference in functional connectivity during naturalistic viewing conditions. Neuroimage 2017;157:521–530.

